# Multimodal analysis of cell-free DNA identifies epigenetic biomarkers for amyotrophic lateral sclerosis diagnosis and progression

**DOI:** 10.64898/2026.03.20.711195

**Authors:** Sebastian Michels, Chaorong Chen, Wolfgang P. Ruf, M. Madhy Garcia Garcia, Frederick J. Arnold, Zhuoxing Wu, Craig L. Bennett, Daniel Shams, Leslie M. Thompson, Alyssa C. Walker, Dennis W. Dickson, Leonard Petrucelli, Johannes Dorst, Mercedes Prudencio, Wei Li, Albert R. La Spada

## Abstract

The role of the epigenome in age-related neurodegenerative disorders remains understudied. Here, we analyzed circulating cell-free DNA (cfDNA) from blood to detect methylation changes as a liquid-biopsy for Amyotrophic Lateral Sclerosis (ALS). Our study included 20 patients with sporadic ALS, 10 patients with C9orf72-associated ALS, 10 asymptomatic carriers of the C9orf72 repeat expansion mutation, and 21 non-disease controls. Following targeted enzymatic methyl-sequencing (EM-seq) of ∼4 million CpG sites, we detected numerous differentially methylated genes, including several implicated in ALS disease risk and pathogenesis. By integrating multiple epigenetic features, we delineated a distinct epigenetic signature, which achieved an average area under the curve (AUC) of 0.91 ± 0.10 upon receiver operator characteristic (ROC) analysis, which enabled detection of ∼70% of ALS patients with close to 100% specificity. Furthermore, we also identified a set of genes whose methylation status significantly correlated with clinical disease progression and cerebrospinal fluid (CSF) neurofilament levels. Our results reveal the potential of cfDNA-based biomarkers to accurately diagnose ALS and potentially predict disease progression.

## Introduction

Amyotrophic Lateral Sclerosis (ALS) is a relentlessly progressive neurodegenerative disease characterized by the rapid loss of motor neurons in the cortex, brainstem, and spinal cord. With an annual incidence of 2 to 4 per 100,000 in the European population, ALS represents the most common motor neuron disease. For the vast majority of patients suffering from ALS, no effective therapy has yet been found to prevent, significantly slow, or halt disease progression. Currently, there are no specific laboratory tests or imaging parameters that can reliably detect ALS. Instead, diagnosis is based on clinical examination indicating upper and lower motor neuron degeneration in combination with the exclusion of another underlying disease. Genetic screening for known ALS-associated mutations is generally recommended; however, these mutations are only found in 70% of familial ALS (fALS) and 15% of sporadic ALS (sALS) cases. Given that up to 10% of all cases are familial, genetic testing alone is insufficient for making an early diagnosis for the majority of ALS patients.

The most established liquid-based biomarkers in ALS are neurofilaments (NfLs), which are structural components of neurons that are released into the CSF and blood from dying myelinated axons. Elevated NfL levels correlate with disease severity and shorter survival in ALS (1). However, their diagnostic value is limited, as increased NfLs are also observed in many other disorders, including inflammatory radiculoneuropathy, stroke, dementia, multiple sclerosis, Huntington’s disease, and COVID-19 infection (2, 3). Currently, biomarker candidates are being developed to detect mis-spliced RNAs or neoepitopes arising from the loss of nuclear TDP-43 function (4, 5). Although these changes can be detected in presymptomatic genetic carriers and ALS patients, their specificity is limited, as TDP-43 pathology is present in several other neurodegenerative disorders and in up to 36% of healthy elderly individuals (6). Consequently, a combination of different biomarkers will likely be necessary to achieve a reliable early ALS diagnosis.

Emerging evidence suggests that epigenetic changes, accumulated over time due to environment and lifestyle factors, contribute to ALS pathogenesis and may influence disease progression. The primary epigenetic mechanism involves the chemical modification of DNA through the methylation of the C5 position of cytosine, forming 5-methylcytosine (5mC). Alterations in DNA methylation, especially in promoter regions, are known to regulate gene expression, either by recruiting proteins involved in gene repression or by inhibiting the binding of transcription factors. Specifically, promoter hypermethylation often acts as a surrogate for gene silencing, while hypomethylation frequently favors increased gene expression. Beyond its essential function in retroviral element silencing, genomic imprinting, and X chromosome inactivation, a stable and unique DNA methylation pattern is required to regulate tissue-specific gene transcription in somatic cells (7). Twin studies (8) and prospective population-based studies (9) indicate an expected heritability of ∼50% in ALS. This incomplete genetic penetrance suggests that additional factors, such as epigenetic mechanisms, likely contribute to ALS risk. Indeed, post-mortem tissue from cortex and spinal cord from ALS patients typically displays changes in DNA methylation affecting immune and inflammatory pathways (10). In addition, a recent blood-based epigenome-wide association study (EWAS) identified differentially methylated genes enriched in immune pathways, energy metabolism, and cholesterol biosynthesis (11). Notably, lower motor neurons of patients with sALS and C9orf72-associated ALS (C9-ALS) exhibit higher methylation levels compared to controls (12).

Recently, the analysis of circulating cell-free DNA (cfDNA) has emerged as a versatile biomarker across various clinical fields based on the concept of a “liquid biopsy”. In oncology, cfDNA enables the sensitive detection of minimal disease activity and treatment response, facilitating personalized treatment strategies for different cancer entities, including lung, breast, pancreatic, and colorectal (13). Notably, one of us successfully implemented a methylation signature derived from plasma-based cfDNA for early detection and monitoring of hepatocellular carcinoma (HCC). The composition of a 77-CpG containing signature demonstrated remarkable specificity (91%) and high sensitivity (76%) in detecting early-stage HCC, surpassing established biomarkers such as serum alpha-fetoprotein (AFP), GALAD score, and ultrasound (14). Additionally, cfDNA can be used to monitor organ rejection (15), and is utilized for non-invasive prenatal testing for genetic abnormalities (16). Circulating cfDNA mainly originates from apoptotic and necrotic cells and is released into the CSF or blood depending upon the tissue of origin, though some cfDNA is found in extracellular vesicles (17). Dying cells typically upregulate nucleases that cleave the cell’s genome into small fragments; hence, nucleosome regions are better protected against enzymatic degradation, whereas open chromatin regions are more susceptible to endonuclease activity (18). Consequently, detectable cfDNA typically exhibits a nucleosome footprint, ranging between 120-220 base pairs. Here we performed a pilot proof-of-concept study to investigate whether sALS or C9-ALS patients exhibit epigenetic signatures capable of discriminating ALS from non-disease controls. We also took advantage of extensive clinical natural history data on these ALS patients to determine if disease-associated epigenetic signatures can predict disease progression. To achieve these goals, we established a multimodal epigenetic sequencing analysis (MESA) model based upon targeted enzymatic methyl-seq (EM-seq) analysis of serum-derived cfDNA from sALS patients, C9-ALS patients, C9-ALS carriers, and non-disease controls. Our results reveal the potential of cfDNA-based biomarkers to accurately diagnose ALS and potentially predict disease progression.

## Results

### ALS patients display a distinct methylome signature

We ascertained a study cohort consisting of 20 patients with sALS, 10 patients with C9-ALS, 10 presymptomatic C9-ALS carriers, and 21 non-disease controls (Table 1), with extensive clinical documentation of disease course and special studies, including measurement of neurofilaments. We isolated cfDNA from their banked serum samples, and after confirming that isolated cfDNA samples exhibit a peak distribution of DNA fragments migrating at ∼170 bp (Figure 1A), we then performed targeted methylome sequencing on all 61 subjects, using the Enzymatic Methyl-seq (EM-seq) method, followed by unbiased target enrichment with the TWIST Human Methylome Panel^®^ of biotinylated probes that provide coverage of ∼4 million CpG sites across the human genome <https://www.twistbioscience.com>. With the exception of one sALS sample, we achieved satisfactory quality control and sufficient unique read depth at ∼45x to permit in-depth epigenetic analysis of the resultant sequencing data for 60 samples, and we detected hundreds of *de novo* differentially methylated regions (DMRs) and associated genes between these four groups (Figure 1, B-E). To assess the importance of achieving ∼45x sequencing depth, we conducted a simulation analysis and found that a substantial portion of DMRs would be missed at sequencing depths less than 45× (Supplemental Figure 1).

**Figure 1.**
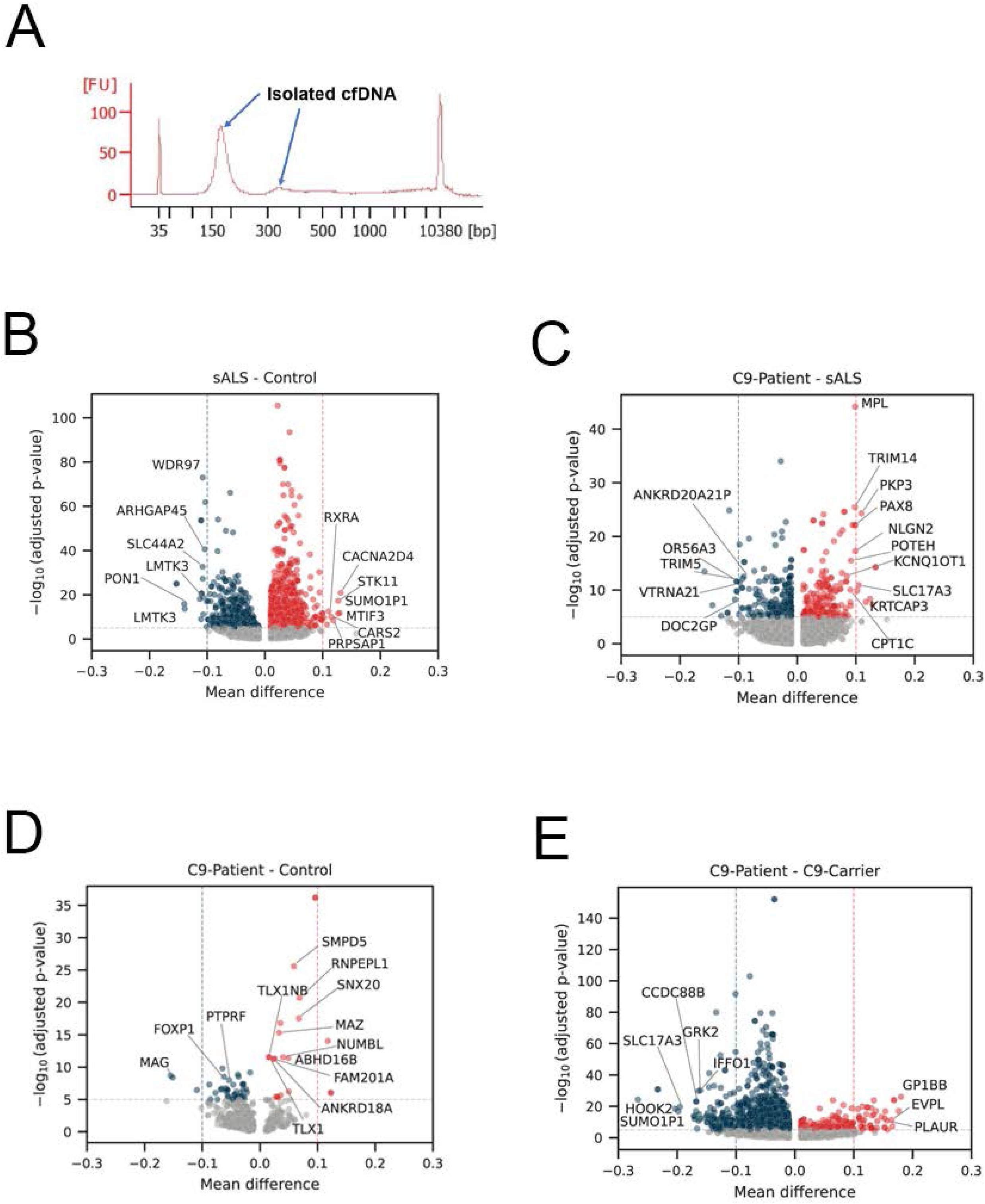
Numerous differentially methylated regions (DMRs) are detectable in plasma cell-free DNA obtained from ALS patients and carriers. **A)** Here we see the size distribution of cell-free DNA obtained from 1.5 mL of blood plasma. Note the major peak at ∼170 bp that perfectly corresponds to the size of DNA protected by a single nucleosome, and a minor peak for a di-nucleosome protected DNA fragment at ∼330 bp. B-E) Volcano plots showing differentially methylated genes as a function of mean methylation difference (x axis) identified by DMR analysis of the following cohorts: **B)** sALS patients vs. non-disease controls; C) C9-ALS patients vs. sALS patients; D) C9-ALS patients vs. non-disease controls; and E) C9 ALS patients vs. C9-ALS carriers. DMRs with an adjusted p-value < 1 × 10^−5^ are colored.

**Table 1.** Patient demographics and Laboratory data.

|  | sALS | C9-ALS | C9-Carrier | Controls |
| --- | --- | --- | --- | --- |
| **N** | 19<sup>A</sup> | 10 | 10 | 21 |
| **Age (IQR) – yr** | 62<br>(53 – 69) | 59.5<br>(55.75 – 62.75) | 44.5<br>(34.25 – 59.75) | 68<br>(35 – 74) |
| **Female sex – no. (%)** | 9 (47.4%) | 3 (30%) | 8 (80%) | 9 (42.9%) |
| **Disease onset** |  |  |  |  |
| Spinal % | 78.9 | 80 | - | - |
| Bulbar % | 21.1 | 20 | - | - |
| **ALS-FRS-R**<br>**(IQR)** | 38<br>(33.5 – 44.0) | 38<br>(31.5 – 40.0) |  |  |
| **Progression Rate<sup>B</sup>**<br>**(IQR)** | 0.80<br>(0.47 – 0.97) | 0.67<br>(0.62 – 1.25) | - | - |
| **pNfH [pg/mL]**<br>**(IQR)** | 3000<br>(1419.5 – 3877.5) | 2202<br>(1134.5 – 4053) |  |  |
| **Concentration cfDNA**<br>**[ng/mL]** | 1.3 ± 2.8 | 5.5 ± 7.4 | 10.4 ± 9.2 | 2.8 ± 2.4 |
| **Total amount cfDNA**<br>**[ng]** | 2.4 ± 5.3 | 8.2 ± 11.1 | 20.7 ± 19.2 | 6.6 ± 6.2 |

sALS = sporadic ALS patients, C9-ALS = patients with a repeat expansion in the C9orf72 gene, C9-Carrier = asymptomatic carrier of a repeat expansion in the C9orf72 gene; IQR = interquartile range; ALS-FRS-R = ALS functional rating scale revised; pNfH = phosphorylated Neurofilament heavy chain. No statistical differences for ALS-FRS-R, disease onset, progression rate, and pNfH levels between sALS and C9-ALS using t-test.
<sup>A</sup>One sALS patient with insufficient cfDNA sequencing reading depth was excluded.
<sup>B</sup>Progression rate was calculated by date of onset to last visit for lost ALS-FRS-R points / months

When we extended our analysis to single CpG resolution, we identified 2,382 CpG sites with significantly different methylation between sALS patients and controls, and 1,801 CpG sites with significantly different methylation between C9-ALS carriers and controls. Clustering of the top 1000 differentially methylated CpG sites (ranked by adjusted p-values) for these cohort comparisons yielded clearly distinguishable methylation patterns (Figure 2, A and B).

**Figure 2.**
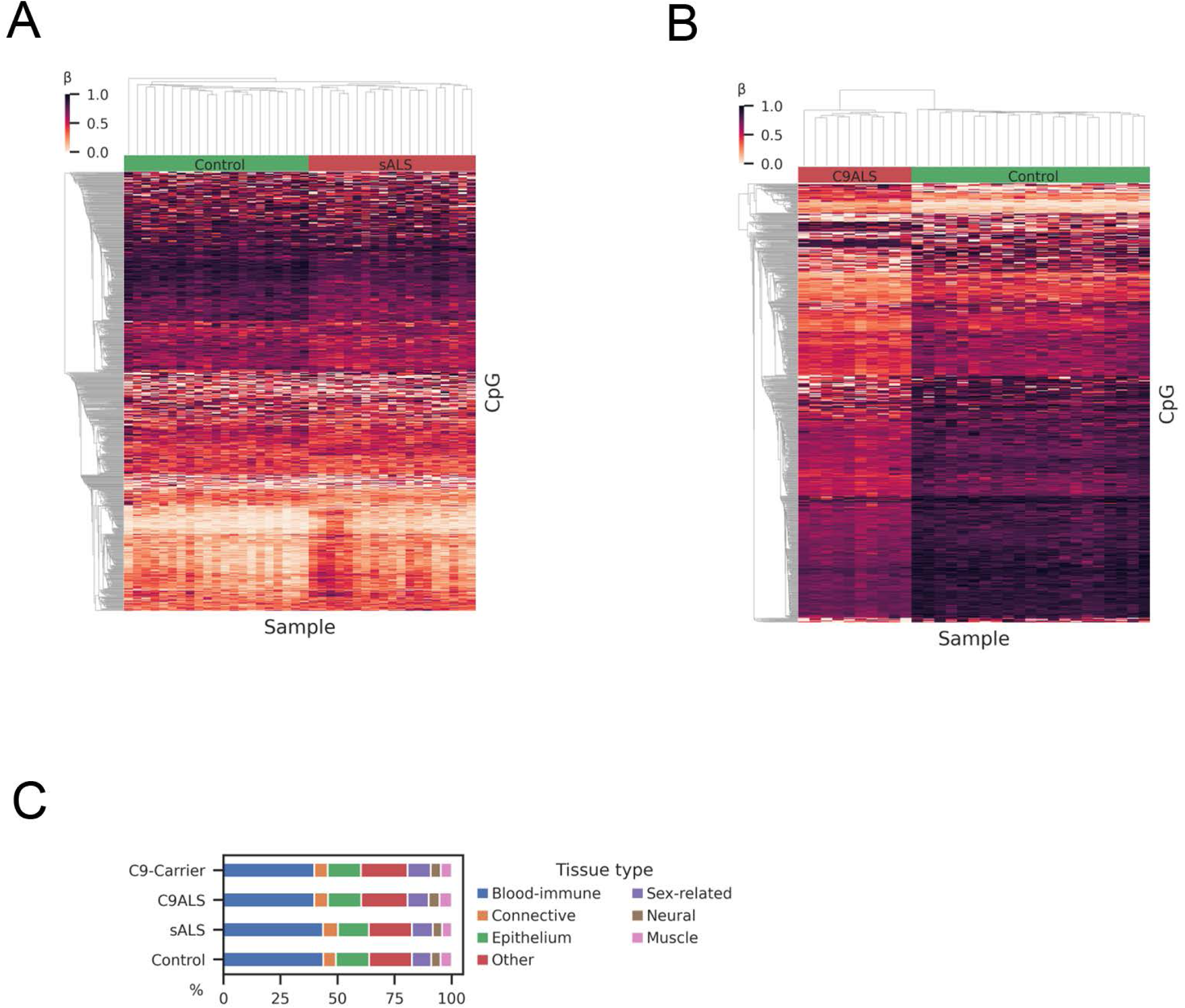
Clustering of differentially methylated CpGs yields clear separation of ALS patient groups from non-disease controls. **A-B**) Clustering of patient groups based upon detection of differentially methylated CpGs in cfDNA samples subjected to targeted EM-seq. Here we clustered the top 1,000 differentially methylated CpGs for: **A**) sALS patients vs. non-disease controls, and **B**) C9-ALS carriers vs. non-disease controls. Note that each column is a single sample, and each row is a single CpG site. The threshold for significantly differentially methylated CpG sites was set at p < 0.001, based upon applying the Mann-Whitney U-test. **C**) Tissue-of-origin results based upon UXM deconvolution analysis of cell-type-specific methylation markers that map to seven different tissue types for each of the cohort groups. Note representation of different tissue types between C9-ALS patients, C9-ALS carriers, sALS patients, and non-disease controls is virtually identical, with ∼4.5% of cfDNA derived from neural tissue for each cohort.

### Deconvolution analysis reveals the origin of cfDNA

To determine the tissue of origin of cfDNA using tissue-specific methylation patterns (19), we applied the UXM algorithm to deconvolute our cfDNA samples into 39 different cell types (20), grouped into seven broad tissue categories: blood immune cells, connective tissue, epithelium, sex-related tissue, neural tissue, muscle, and other. Our results revealed that ∼4.5% of cfDNA is derived from neural tissue (Figure 2C & Supplemental Table 1), suggesting that a blood-based liquid biopsy can reliably detect methylation changes originating in the central nervous system (CNS). To better understand the origin of the neural cfDNA signal, we performed an expanded tissue deconvolution analysis incorporating brain cell subtypes (21), and determined that excitatory neurons are predominant contributors to neural-derived cfDNA (Supplemental Figure 2). Furthermore, our analysis of cfDNA tissue type did not reveal significant global differences in cfDNA tissue-of-origin among these cohorts. To assure accuracy of the deconvolution, we performed a simulation study using *in silico* mixtures of sequenced reads. For 50 mixtures at 5 different neuron sample concentrations varying from 0 to 10%, the simulation results confirmed that the UXM algorithm can accurately detect cfDNA tissue-of-origin (Supplemental Figure 3).

### ALS patients show cfDNA methylation alterations in disease-relevant genes

The most established biological mechanism for epigenetic gene regulation is the methylation status of the promoter region. While promoter hypermethylation typically leads to decreased gene expression, promoter hypomethylation results in increased gene expression. To further explore the relevance of differentially methylated genes in cfDNA, we applied a stringent filter to differentially methylated regions (DMRs) in the promoter region. Specifically, we retained promoter-overlapping DMRs with ≥ 10 CpG sites, an adjusted p-value < 1 × 10^−5^ between groups, and a mean methylation difference greater than or equivalent to ± 0.1 between groups. DMRs in the X-chromosome were excluded, as they naturally exhibit significant differences due to X-inactivation between the male and female sex. Unbiased *de novo* DMR analysis of sALS patients vs. non-disease controls revealed 18 genes with significant promoter methylation changes (Table 2). Of these highly significant hits, the DMR associated with *PON1*, the gene encoding Paraoxonase 1, exhibited the greatest change in methylation pattern. In the sALS cohort, the *PON1* DMR was hypomethylated with an average promoter methylation of 43.6% compared to 57.6% in controls (p = 5.0 x 10^−18^). *PON1* encodes an enzyme found in high-density lipoprotein (HDL) (22). While the association of ALS and cardiovascular disease is ambiguous, there is evidence that alterations in cholesterol metabolism are a risk factor and a negative prognostic factor in ALS (23, 24). Furthermore, several genetic single nucleotide polymorphism (SNP) coding variants in *PON1* have been associated with increased susceptibility to sALS, including Q192R, L55M, and A162G (25–28), and another independent study reported a common haplotype in the *PON1* promoter that is significantly linked to sALS disease risk (29).

**Table 2:** Top Hits for Promoter Differentially Methylated Regions for sALS Patients vs Controls.

| Gene | Methylation in sALS (%) | Methylation in controls (%) | Mean methylation difference | p-value (MWU) |
| --- | --- | --- | --- | --- |
| PON1 | 43.59 | 57.57 | -13.98 | 5.04E-18 |
| LMTK3 | 42.90 | 56.78 | -13.88 | 1.12E-15 |
| SLC17A3 | 39.03 | 50.10 | -11.07 | 1.20E-10 |
| SLC44A2 | 51.89 | 62.69 | -10.79 | 1.54E-35 |
| WDR97 | 62.31 | 73.08 | -10.77 | 9.89E-77 |
| SMIM24 | 50.74 | 61.42 | -10.68 | 5.06E-13 |
| ARHGAP45 | 69.86 | 80.28 | -10.42 | 1.13E-43 |
| MPL | 56.63 | 66.96 | -10.33 | 2.90E-65 |
| GLB1L | 40.99 | 51.06 | -10.07 | 4.12E-19 |
| SEPTIN9 | 39.13 | 28.92 | 10.21 | 1.73E-11 |
| DTX2 | 58.74 | 47.99 | 10.75 | 4.75E-08 |
| RXRA | 36.36 | 25.30 | 11.05 | 1.64E-14 |
| CARS2 | 29.81 | 18.36 | 11.45 | 6.73E-12 |
| PRPSAP1 | 45.25 | 33.40 | 11.85 | 5.90E-10 |
| MTIF3 | 39.65 | 26.98 | 12.67 | 2.61E-13 |
| STK11 | 28.28 | 15.58 | 12.70 | 1.64E-19 |
| SUMO1P1 | 41.05 | 28.11 | 12.93 | 1.00E-13 |
| CACNA2D4 | 25.85 | 12.73 | 13.12 | 2.57E-23 |

sALS = sporadic Amyotrophic Lateral Sclerosis, MWU = Mann-Whitney-U-Test; genes are ranked by mean methylation difference. Non-coding genes are excluded.

Other ALS disease-relevant genes were also identified as displaying markedly different methylation patterns in promoter DMRs, including the *LMTK3* gene that encodes the lemur tail kinase-3 protein and the *STK11* gene that encodes the protein LKB1, a well-known tumor suppressor (Table 2). LMTK3 belongs to a family of lemur tyrosine kinases that are highly expressed in layer II to VI of the motor cortex and in the CA1-3 area in the dentate gyrus of the hippocampus (30). While LMTK3 normally functions in neurite extension and apoptosis, axonal transport, and endosomal vesicle trafficking (31), (32), LMTK3 can regulate cortical excitability through its interactions with the chloride-potassium transporter KCC2 (33), potentially implicating LMTK3 in excitotoxicity, which is a defining feature of ALS. Notably, post-mortem studies of sALS patient frontal cortex revealed increased expression of LMTK3 in neuron axons and dendrites (34), which could be explained by hypomethylation of the *LMTK3* promoter DMR observed in our sALS patient cohort and found to be also present in C9-ALS presymptomatic carriers (Table 2). In the case of *STK11*, an unbiased integrative multi-omics study of sALS found that copy number variants (CNVs) in *STK11* are associated with ALS disease risk and might represent a potential biomarker for sALS patient stratification (35), although the effect of the CNV increase at the *STK11* locus upon gene expression was not determined. Whether hypermethylation of the *STK11* promoter DMR detected by our methylome sequencing analysis of sALS patients is related to the CNV increase observed at this locus in sALS patients deserves consideration.

Since promoter methylation plays an important role in regulating gene expression, we further estimated the methylation level of promoter DMRs for ALS-associated genes (Figure 3). For *TARDBP*, which encodes the TDP-43 protein, we noted significant hypomethylation in both sALS patients and C9-ALS patients compared to non-disease controls, but remarkably we observed no differences in hypomethylation between C9-ALS carriers and non-disease controls (Figure 3), suggesting that epigenetic changes in the TDP-43 gene do not occur until after initiation of clinically significant disease. Interestingly, a post-mortem analysis of ALS patient frontal cortex found that *TARDBP* hypomethylation correlates with age of disease onset (36). Analysis of the *C9orf72* gene revealed marked variability in methylation status, with markedly increased promoter methylation in C9-ALS patients and C9-ALS carriers (Figure 3), in agreement with prior studies (37). While there were trends towards hypomethylation of *SOD1* and *AGRN* in C9-ALS patients and carriers, and evidence for hypermethylation of *FUS* in C9-ALS patients, these differences did not achieve statistical significance (Figure 3). We also noted marked promoter hypomethylation of the *ATXN2* gene in C9-ALS patients, especially when compared to non-disease controls (Figure 3). These results indicate that ALS-linked genes are subject to CpG-level epigenetic alterations that can be detected in cfDNA isolated from serum of sALS and C9-ALS patients.

**Figure 3.**
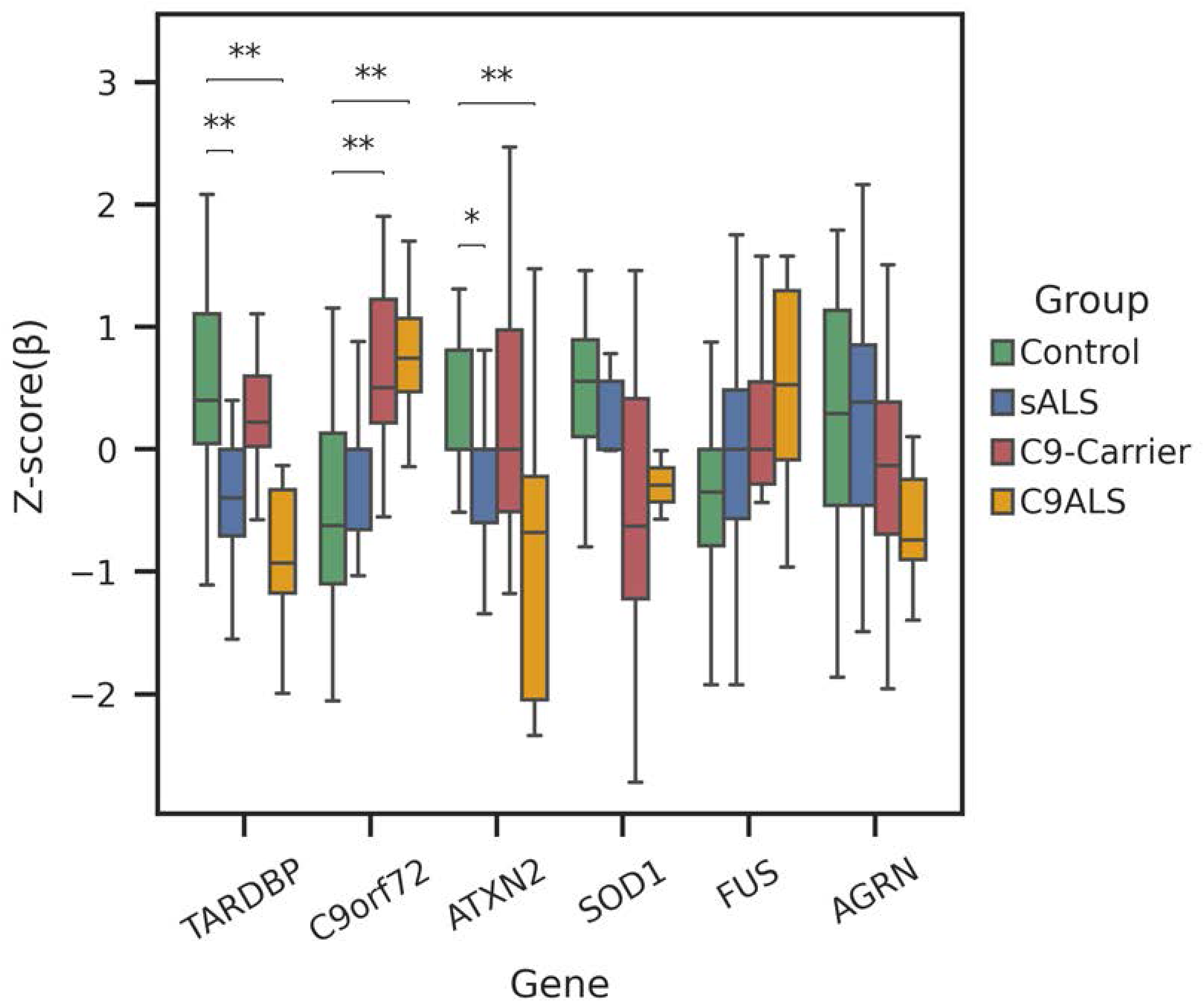
CpG methylation levels in promoters of certain ALS-associated genes vary significantly between sALS patients, C9-ALS patients, C9-ALS carriers, and non-disease controls. Here we see a plot of the methylation levels for the top promoter DMCs of the listed ALS-associated genes measured in cfDNA samples obtained from blood plasma after normalization. The Y-axis corresponds to the methylation level, which is given as a Z-score. For this comparison, a positive Z-score means hypermethylation, while a negative Z-score equates to hypomethylation. Statistical comparisons for significant differences are shown based upon applying the Mann-Whitney U-test. *p < 0.05, **p < 0.01. Error bars = s.d.

### Multimodal epigenetic sequencing analysis of cfDNA enables detection of ALS patients

To permit a more comprehensive analysis of epigenetic signatures in cfDNA samples from ALS patients, we extracted three additional epigenetic features from our methylome sequencing data: 1) DNase I hypersensitive site (DHS) region methylation, 2) nucleosome occupancy, and 3) window protection score. DHSs are established markers of regulatory DNA, which reveal novel relationships between chromatin accessibility, transcription and DNA methylation (38); nucleosome occupancy reflects the likelihood that a given base pair in a cell will be occupied by a nucleosome; and window protection score is a measure of the dynamics of DNA fragmentation as it relates to nucleosome positioning (18). We have previously combined these three metrics with differential CpG site methylation to build a machine learning model for colorectal cancer detection, which we called multimodal epigenetic sequencing analysis (MESA) (39), to identify distinct epigenetic features for evaluation of methylome sequencing data from cfDNA. To determine if cfDNA-based epigenetic properties can distinguish ALS patients from non-disease controls, we applied the MESA pipeline to our data set and trained a model on the most significant markers for these four modalities. To test the performance of MESA in detecting sALS patients, we used leave-one-out cross-validation (LOOCV) and obtained an average area under the curve (AUC) of 0.91 in the receiver operator curve (ROC) characteristic analysis (Figure 4). Notably, with a false positive rate of 0% (i.e. 100% specificity), comparison of cfDNA epigenetic signatures enabled us to detect 70% of sALS patients. To determine the robustness and generalizability of MESA, we considered a recently published cfDNA whole-genome bisulfite sequencing data set obtained from 10 Caucasian ALS patients and 8 non-disease controls (40). When we applied our MESA algorithm to this independent cohort, we obtained an AUC of 0.85, cross-validating its strong performance for different sequencing methods and geographically diverse cohorts. We also inspected inter-individual biomarker variability for our top 100 differentially methylated CpG (DMC) sites, quantified by coefficient of variation, and noted that methylation variability was much greater in C9-ALS carriers and sALS patients compared to non-disease controls (Supplemental Table 2). This variability indicates that cfDNA methylation is capturing biological differences between different cohorts and shows that cfDNA-based biomarkers exhibit greater variability in diseases groups.

**Figure 4.**
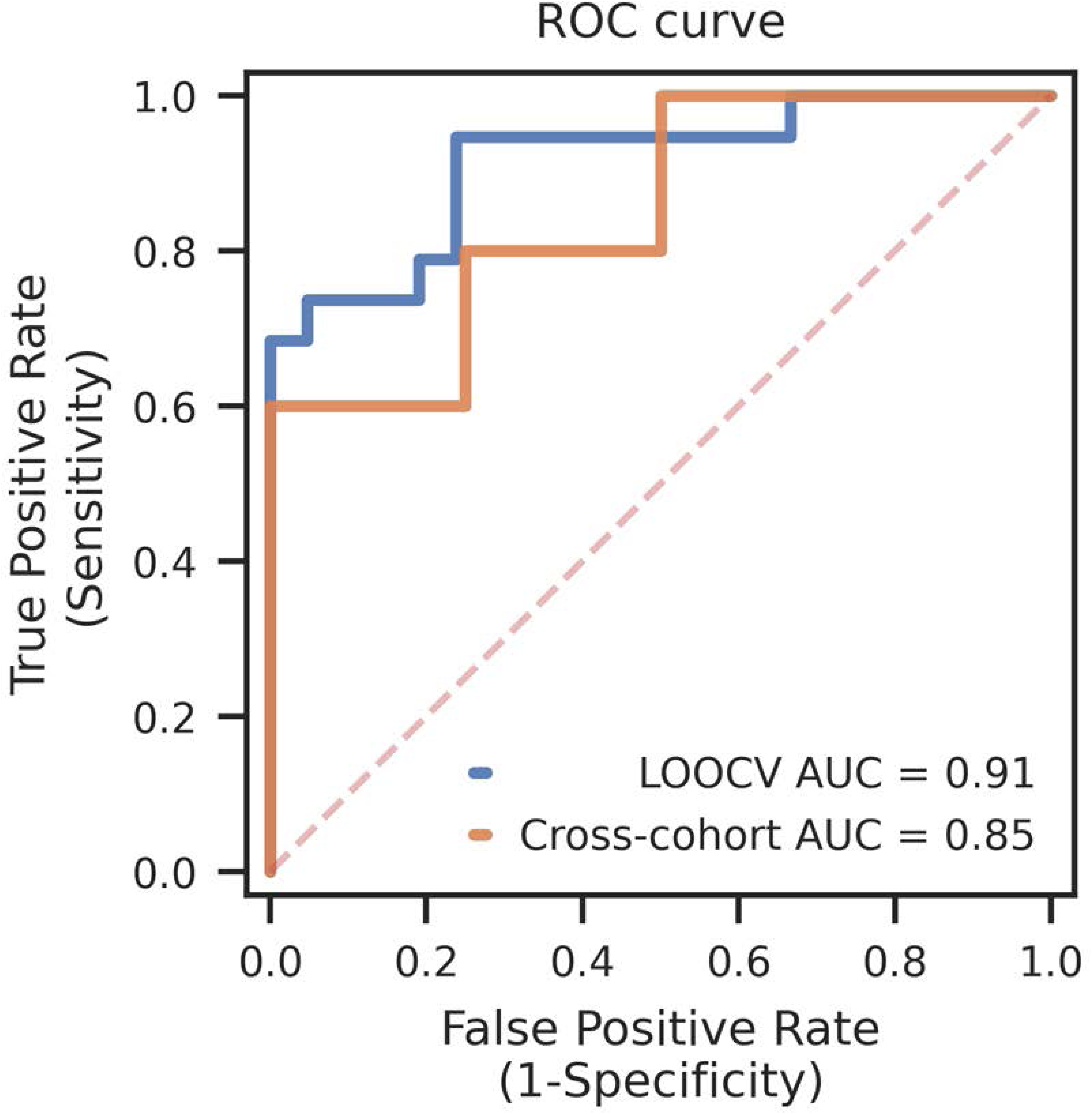
MESA analysis can accurately detect sALS patients in comparison to non-disease controls. Using MESA, we performed receiver operator characteristic curve analysis for the most significantly different cfDNA multimodal epigenetic markers for sALS patients (n = 19) vs. non-disease controls (n = 21). LOOCV analysis yielded an area under the curve (AUC) of 0.91, and was then applied to an independent cross-validation cohort (Cross-cohort) of sALS patients (n = 10) vs. non-disease controls (n=8), yielding an AUC of 0.85.

### CSF neuroAilament levels and differential promoter methylation correlate with ALS disease progression

At present, there is an urgent need for reliable biomarkers to track disease progression in ALS patients. The measurement of neurofilaments is being utilized as a biomarker in ALS, as neurofilaments may reflect axonal damage and ongoing motor neuron degeneration. We thus considered the utility of neurofilament levels as a predictor of disease progression in the 18 sALS patients and seven C9-ALS patients from our study cohort for which neurofilament levels in blood and CSF had been obtained and for whom monthly longitudinal clinical assessments had been performed. When we compared disease progression between these 18 sALS patients and seven C9-ALS patients, we noted more rapid disease progression in sALS patients, based upon a higher median loss of ALS-FRS-R points per month, as sALS patients declined at an average rate of 0.80 ALS-FRS-R points per month, while C9-ALS patients declined at an average rate of 0.67 ALS-FRS-R points per month, but this difference did not achieve statistical significance due to the small cohort sizes. In line with previous studies (41), the level of phosphorylated neurofilament heavy chain (pNfH) in CSF from our 18 sALS and seven C9-ALS patients correlated with this difference in disease progression, with sALS patients displayed higher median CSF pNfH levels in comparison to C9-ALS patients (3,000 pg/mL vs. 2,202 pg/mL). However, these differences were not statistically significant owing to the small cohort sizes. The sALS cohort and C9-ALS cohort were similar in terms of location of disease onset (∼80% spinal and ∼20% bulbar; p = 0.949), age (p = 0.189), and sex (p = 0.135). When we evaluated CSF pNfH as a biomarker for ALS disease progression in all 25 ALS patients, we observed a significant correlation between the slope of ALS-FRS-R decline and the increase in pNfH levels (Figure 5A). This data supports pNfH levels as an indicator of disease progression in our study cohort.

**Figure 5.**
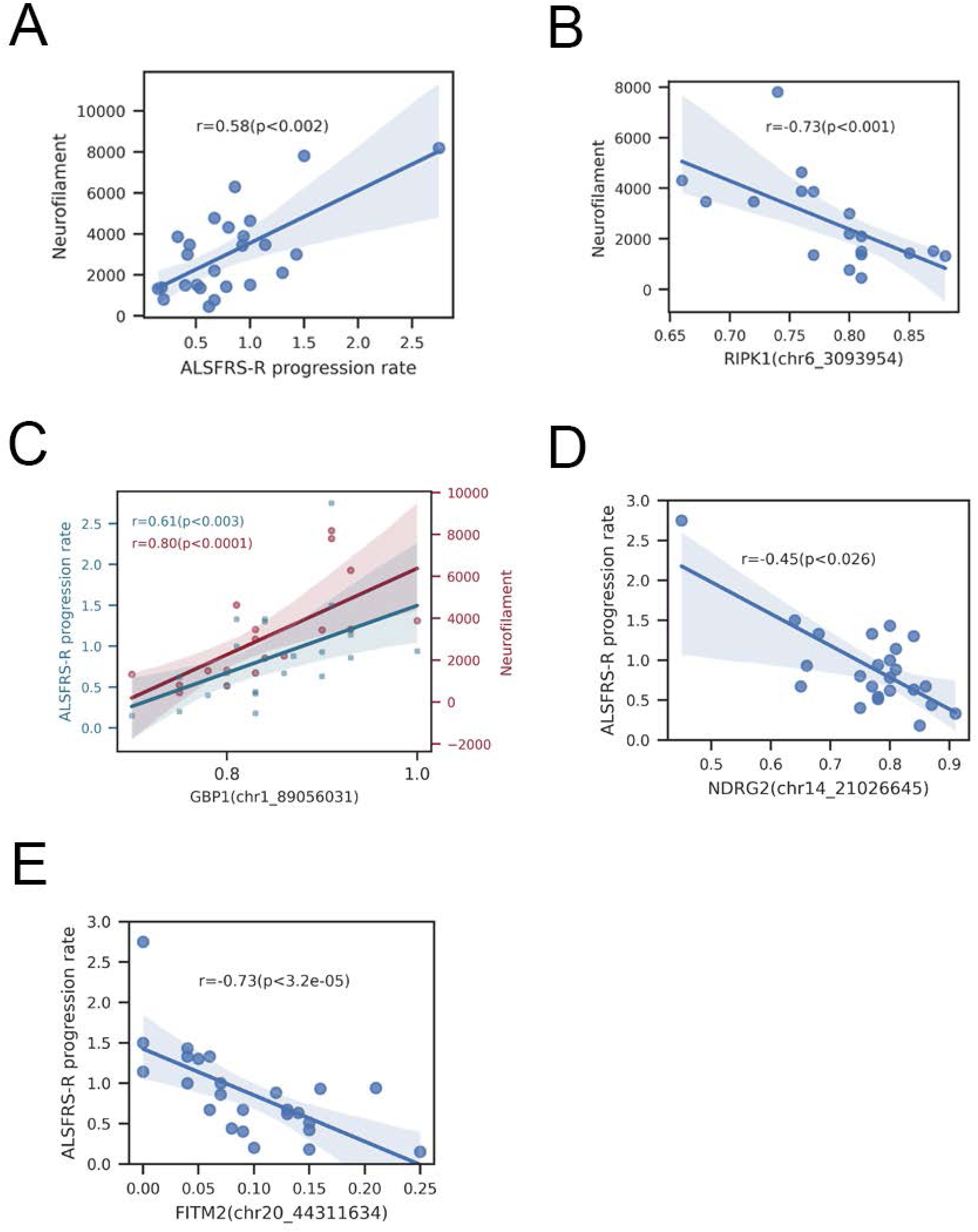
NeuroIilament levels and disease progression rates correlate with single gene methylation changes in ALS patients. **A)** Plot of phosphorylated neurofilament heavy chain (pNfH) levels as a function of ALS-FRS-R rate of disease progression for 25 ALS patients (sALS, n = 18; C9 ALS, n = 7). Statistical comparison is based upon Spearman rank correlation analysis. **B)** Plot of phosphorylated neurofilament heavy chain (pNfH) levels as a function of *RIPK1* gene methylation for sALS patients (n = 18). Statistical comparison is based upon Spearman rank correlation analysis. **C)** Plot of both phosphorylated neurofilament heavy chain (pNfH) levels and ALS-FRS-R rate of disease progression as a function of *GBP1* gene methylation for sALS patients (n = 18). Statistical comparison is based upon Spearman rank correlation analysis. **D)** Plot of ALS-FRS-R rate of disease progression as a function of *NDRG2* gene methylation for sALS patients (n = 19). Statistical comparison is based upon Spearman rank correlation analysis. **E)** Plot of ALS-FRS-R rate of disease progression as a function of *FITM2* gene methylation for sALS patients (n = 19). Statistical comparison is based upon Spearman rank correlation analysis.

We next tested if pNfH levels are correlated with promoter CpG site methylation differences in selected genes. Among genes with the most significant DMRs, we identified a negative correlation between pNfH CSF levels and methylation levels of the *RIPK1* gene in sALS patients (Figure 5B & Table 3). *RIPK1* encodes the receptor-interacting serine/threonine protein kinase 1, is upregulated in post-mortem spinal cord tissue from ALS patients (42), and its serum concentrations were found to be higher in ALS patients (43). In contrast, in the case of the *GBP1* gene, we detected a positive correlation between pNfH CSF levels, and *GBP1* methylation (Figure 5C & Table 3). *GBP1* encodes a large GTPase involved in cellular signal transduction responses to inflammation and environmental stressors.

**Table 3:** Top 10 Differentially Methylated Genes correlating with pNfH CSF values.

| Gene | Spearman correlation with pNfH | Spearman (pNfH) p-value | Spearman correlation with ALS-FRS-R slope | Spearman (ALS-FRS-R) p-value |
| --- | --- | --- | --- | --- |
| GBP1 | 0.80 | 5.8E-05 | 0.61 | 2.6E-03 |
| NPHP3 | 0.75 | 1.5E-04 | 0.26 | 0.22 |
| RIPK1 | -0.73 | 6.2E-04 | -0.23 | 0.31 |
| EHMT1 | -0.71 | 8.9E-04 | -0.20 | 0.38 |
| ZNF528 | -0.71 | 1.4E-04 | -0.35 | 0.12 |
| ULK1 | 0.70 | 1.4E-04 | 0.45 | 0.02 |
| OFCC1 | 0.67 | 3.1E-03 | 0.49 | 0.03 |
| MTHFD2 | 0.67 | 3.4E-03 | 0.06 | 0.81 |
| PC | 0.66 | 7.9E-04 | 0.36 | 0.07 |
| ZNF391 | 0.66 | 5.4E-03 | 0.22 | 0.34 |

pNfH = phosphorylated neurofilament heavy chain; non-coding genes were excluded.

To further determine if CpG methylation differences in a single gene could be a significant predictor of ALS clinical disease progression, we next considered our top promoter differentially methylated CpG (DMC) hits. Interestingly, methylation of a *GBP1* DMC also directly correlated with sALS clinical disease progression (Figure 5C & Table 4). We further noted that DMC methylation of the N-myc downstream-regulated gene 2 (*NDRG2*) exhibited a strong negative correlation with ALS-FRS-R slope (Figure 5D & Table 4), as sALS patients defined as displaying faster disease progression have greatly reduced *NDRG2* promoter CpG site methylation. The NDRG2 protein is known for its role as a tumor-suppressor, inhibiting several key cell pathways, including Akt, NF-κB, and TGF-β signaling (44), but NDRG2 protein is also highly expressed in motor neurons (45), and can promote neurite outgrowth (46) and regulate oxidative stress. Interestingly, splicing dysregulation of the *NDRG2* gene occurs upon knockdown of either FUS/TLS or TDP-43 (47), and increased NDRG2 protein expression, as would be expected upon promoter hypomethylation, has been implicated in disease pathogenesis in ALS model mice (45). Finally, we noted that DMC methylation of *FITM2*, which encodes the fat storage-inducing transmembrane protein 2, displayed a strong negative correlation with clinical ALS disease progression (Figure 5E & Table 4). FIT proteins are critical for promoting the formation of lipid droplets within the endoplasmic reticulum. FITM2 is highly expressed in adipocytes, where it facilitates production of large lipid droplets essential for energy storage and metabolic homeostasis (48). Dysregulation of lipid droplets have been implicated in ALS disease pathogenesis, as TDP-43 loss of function results in metabolic disturbances, including an accumulation of lipid droplets (49). Furthermore, dramatically increased lipid droplet accumulation was detected in skeletal muscle biopsies obtained from fALS patients carrying mutations in *FUS* (50).

**Table 4:** Top 10 Differentially Methylated Genes correlating with ALS Disease Progression.

| Gene | Spearman correlation with pNfH | Spearman (pNfH) p-value | Spearman correlation with ALS-FRS-R slope | Spearman (ALS-FRS-R) p-value |
| --- | --- | --- | --- | --- |
| FITM2 | -0.42 | 0.06 | -0.73 | 3.2E-05 |
| DNAH12 | 0.29 | 0.26 | 0.68 | 1.0E-03 |
| NEBL | 0.41 | 0.09 | 0.67 | 9.3E-04 |
| SLC35A1 | -0.24 | 0.31 | -0.66 | 5.9E-04 |
| HOXA9 | 0.44 | 0.03 | 0.65 | 1.2E-04 |
| SHC2 | -0.12 | 0.61 | -0.64 | 5.4E-04 |
| SIGIRR | -0.32 | 0.12 | -0.64 | 1.9E-04 |
| KDM5B | 0.54 | 0.02 | 0.63 | 2.2E-03 |
| APEH | -0.45 | 0.02 | -0.63 | 2.5E-04 |
| CTIF | 0.59 | 0.02 | 0.63 | 3.0E-03 |

pNfH = phosphorylated neurofilament heavy chain; non-coding genes were excluded.

### Studies of ALS-associated *RIPK1* hypomethylation conAirm expression increase in human patients and implicate RIPK1 expression increase in microglia dysfunction

To determine if highly significant cfDNA promoter methylation differences found to correlate with ALS patient CSF neurofilament burden or disease severity result in altered expression of the implicated genes, we selected *NDRG2* and *RIPK1* for further analysis, as these two target genes are known to be dysregulated in ALS model mice and in human patients (42,43,45). When we measured the RNA expression levels of *NDRG2* and *RIPK1* in the frontal cortex of patients with frontotemporal dementia associated with TDP-43 (FTD) with or without ALS motor neuron disease, we noted increased expression of both these genes in FTD-TDP and FTD-TDP/ALS patients, though the increase only constituted a trend for *NDRG2* (Figure 6A). A previous single-nucleus RNA sequencing study on sALS spinal cord showed that increased expression of *RIPK1* promoted in glial cells is associated with neuroinflammation (51). To validate whether increased *RIPK1* expression leads to inflammation in microglia, we transduced induced microglia-like cells (iTF-microglia), derived from induced pluripotent stem cells (52), with a *RIPK1* lentivirus and stimulated inflammation by treating with lipopolysaccharide (LPS). We observed a robust cytokine response, which was significantly augmented by increased expression of *RIPK1* (Figure 6B). We then assayed the phagocytosis capacity of iTF-microglia subjected to LPS treatment and *RIPK1* expression increase, and we found that increased RIPK1 significantly impaired phagocytosis (Figure 6C), indicating that RIPK1 diminished this neuroprotective function. Another key microglia task is migration, which is involved in the response to injury. To assay microglia migration, which is accentuated when microglia are reactive, we treated iTF-microglia with LPS in the presence or absence of RIPK1, and we observed markedly increased iTF-microglia migration with RIPK1 overexpression (Figure 6D). Furthermore, higher RIPK1 expression led to enhanced cell death (Figure 6E). Overall, these findings support a role for increased RIPK1 expression in promoting glial cell dysfunction and neuroinflammation.

**Figure 6.**
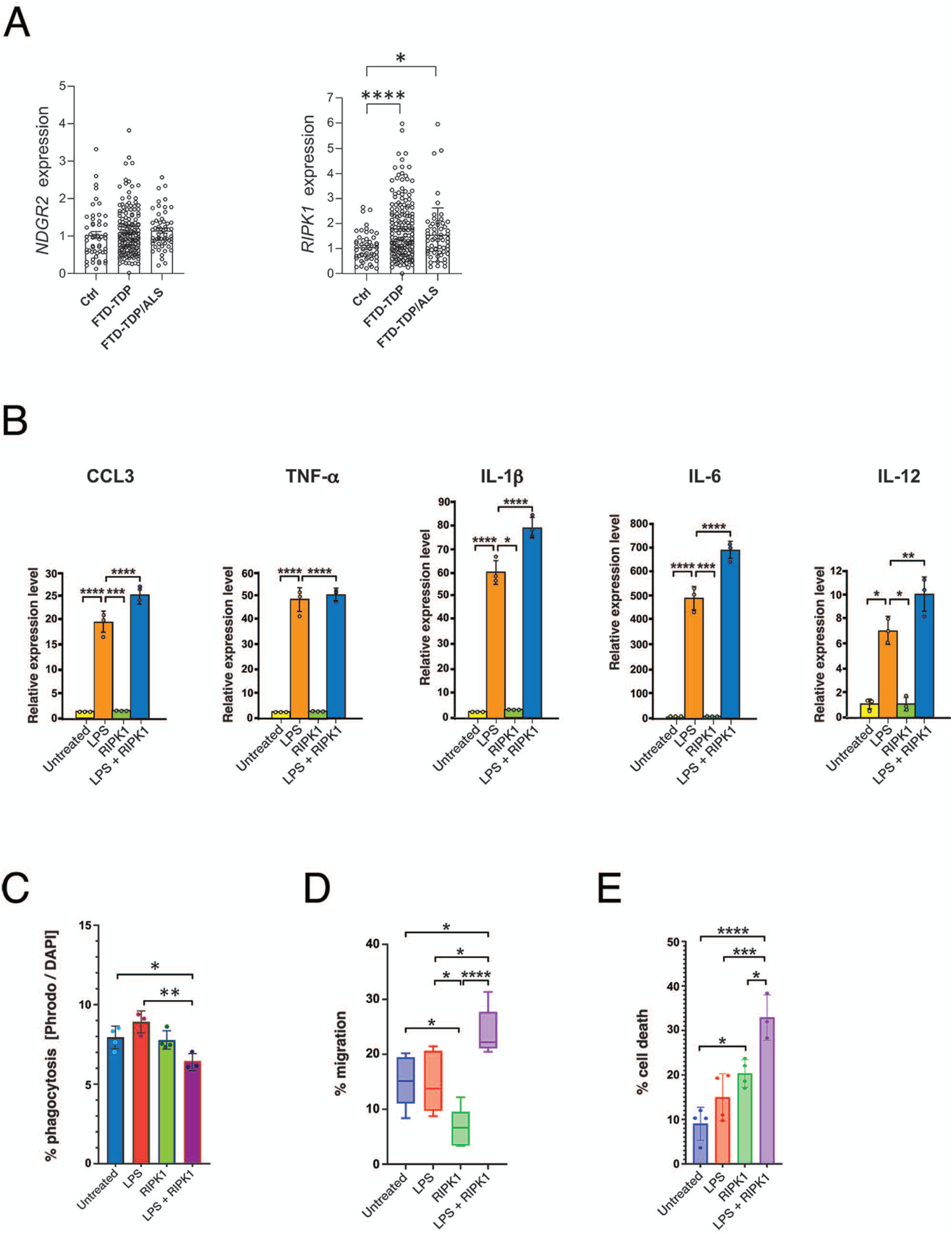
Expression levels of differentially methylated genes in post-mortem FTD-ALS patient cortex and evaluation of effect of RIPK1 expression increase on microglia function. **A)** Quantitative real-time RT-PCR analysis of RNAs isolated from the frontal cortex of FTD patients (n = 167) or FTD-ALS patients (n = 59) with evidence of TDP-43 histopathology on post-mortem examination, as well as non-disease controls (n = 51) to measure the expression level of *NDGR2* and *RIPK1*. Statistical significance was calculated using the Kruskal-Wallis non-parametric test. *p < 0.05, ****p < 0.0001. **B)** iTF-microglia were transduced with a *RIPK1* lentivirus expression or empty vector, and then left untreated or subjected to LPS treatment. We then isolated RNAs and performed qRT-PCR analysis of the indicated cytokine genes. *p<0.05, **p<0.01, ***p<0.001, ****p<0.001; ANOVA with post-hoc Tukey test, n = 3 biological replicates. **C)** iTF-microglia were transduced with a *RIPK1* lentivirus expression or empty vector, and then left untreated or subjected to LPS treatment. We then incubated iTF-microglia with pHrodo™ Red Zymosan BioParticles™ Conjugate, and measured pHrodo™ Red Zymosan BioParticles fluorescence as a ratio of DAPI fluorescence. *p<0.05, **p<0.01; ANOVA with post-hoc Tukey test, n = 4 biological replicates. **D)** iTF-microglia were transduced with a *RIPK1* lentivirus expression or empty vector, and then left untreated or subjected to LPS treatment. To measure migration, iTF-Microglia were plated onto PDL and ECM coated 8 μm transwells, ADP was added as the attractant, and numbers of migrating microglia were counted, *p<0.05, ****p<0.001; ANOVA with post-hoc Tukey test, n = 6 biological replicates. Boxes indicate the 25^th^ percentile to 75^th^ percentile with respective medians marked by a line within the box, while whiskers encompass the full range of scores for each condition. Please see the supporting raw values file for exact scores for each condition. **E)** iTF-microglia were transduced with a *RIPK1* lentivirus expression or empty vector, and then left untreated or subjected to LPS treatment. We then stained iTF-microglia with DAPI and counted the numbers of dead cells based upon amoeboid and blebbing appearance. *p<0.05, ***p<0.001, ****p<0.0001; ANOVA with post-hoc Tukey test, n = 4 biological replicates. Error bars = s.e.m.

## Discussion

Differentiating ALS from other conditions with similar clinical presentations remains a significant diagnostic challenge, particularly in the early stages of the disease. The lack of specific tests and common misinterpretation of ALS-mimicking syndromes pose a substantial burden for patients, especially given the short median life expectancy of about three years. This often leads to prolonged periods of diagnostic uncertainty, unnecessary medical procedures, and delayed enrollment in clinical trials. This delayed diagnosis may close the therapeutic window, such that disease has progressed beyond the point at which disease progression may be attenuated. Over 90% of ALS cases are sporadic and likely to involve genetic susceptibility to environmental risk factors that contribute to the disease through epigenetic changes (10). While there are various methods to gauge epigenetic alteration, an exciting development has been the analysis of cell-free DNA (cfDNA) as a marker of epigenetic change.

Unlike a previous report that restricted analysis to pre-selected targets (53), here we pursued a completely unbiased approach to the analysis of epigenetic change in ALS by performing EM-seq combined with target enrichment via a methylome panel of biotinylated probes that provide coverage of ∼4 million CpG sites across the human genome. Despite our relatively small cohort sizes, we were able to detect hundreds of significantly altered CpG methylation events in both sALS and C9orf72 ALS patients in comparison to non-disease controls. Because of the robustness of our sequencing approach, which offers numerous advantages over sodium bisulfite sequencing that yields much lower complexity sequencing libraries, we could extract other key epigenetic features, including DNase I hypersensitive site methylation, nucleosome occupancy, and window protection score. When we analyzed the top markers for these four features using our MESA machine learning algorithm and performed ROC characteristic analysis, we could readily distinguish the majority of sALS patients (70%) from non-disease controls with 100% specificity, reflecting the obtained AUC of 0.91. We also applied MESA to an independent cohort of 10 sALS patients and 8 controls, and obtained an AUC of 0.85, convincingly demonstrating its broad utility. Given the small size of these respective cohorts, this level of detection bodes well for the application of this strategy as a tool for facilitating ALS disease diagnosis.

Freely circulating cfDNA levels in the bloodstream are physiologically determined by the rate of DNA release and the activity of clearing enzymes such as DNase I, factor VII-activating protease, and factor H (54). For disease conditions characterized by excessive cell death, such as cancer, cfDNA can accumulate to very high levels due to an increased cfDNA release, which subsequently overloads the clearance system. Similarly, due to the aggressive nature of ALS, we anticipated that this rapidly progressive motor neuron degeneration would lead to an enrichment of cfDNA levels originating from neurons and muscle cells. While a higher abundance of muscle-derived cfDNA in ALS has been shown using bisulfite sequencing (55), our results did not indicate alterations in cfDNA composition in ALS patients compared to controls, which may arise from differences in cfDNA isolation and sequencing methods. Compared to bisulfite-sequencing, EM-seq is non-destructive to the input DNA resulting in longer DNA fragments (>10 kb) and better coverage of the methylome allowing the detection of subtle methylation changes (56), which enhances the accuracy of tissue deconvolution. Our overall cfDNA tissue distribution is similar to reports in healthy controls using whole-genome bisulfite sequencing (57). We also did not find that ALS patient samples contained more cfDNA than controls, contrary to a recently published study (55).

Our deep ∼45x methylation sequencing is critical for identifying neuron-derived ALS biomarkers, as it provides the statistical power needed to detect subtle methylation changes between clinical samples, particularly given the relatively low abundance (∼4.5%) of neuronal-derived cfDNA. It is intriguing that our analysis revealed several top methylation changes in the promoter regions of genes that are associated with ALS or centrally implicated in ALS disease pathogenesis. For the TDP-43 gene *TARDBP*, demethylation in the 3’ untranslated region in the motor cortex has previously been associated with the age of ALS onset (36). Our methylation sequencing analysis of cfDNA revealed significant promoter hypomethylation in *TARDBP* in both sALS patients and C9-ALS patients compared to non-disease controls. We also observed significant methylation changes in ALS patient cfDNAs for the promoters of three genes (*PON1, LMTK3*, and *STK11*), whose gene products have all been linked to ALS, either as disease risk associations in the case of *PON1* and *STK11*, or as undergoing marked changes in gene expression (*LMTK3)* in pathologically relevant regions of the CNS in post-mortem studies of ALS patients. These findings indicate that methylation sequencing of ALS patient cfDNA may reflect ongoing cellular processes tied to ALS disease pathogenesis. When we tried to better understand the origin of these neuron-related methylation changes with CellDMC analysis (58), we did not identify any specific cell type as the primary driver. We speculate that the biomarkers detected in cfDNA do not originate from a single cell or tissue type, but instead likely reflect a systemic, whole-body effect involving multiple biological sources, including diseased neurons.

Although the application of cfDNA epigenetic signature analysis as an assay for facilitating ALS disease diagnosis is in and of itself an exciting advance, we also considered use of cfDNA epigenome changes as a biomarker for tracking ALS disease progression. To maximize the likelihood of identifying a correlation between cfDNA epigenetic alterations and ALS disease progression, we focused on genes with the greatest extent of methylation change, selecting our top DMCs. With this approach, we found significant correlations between ALS disease progression and methylation for three genes, *GBP1*, *NDRG2* and *FITM2*. We noted that increased methylation of the *GBP1* gene correlated with an increased rate of ALS disease progression, while decreased methylation of the *NDRG2* and *FITM2* genes correlated with an increased rate of ALS disease progression. As increased expression of the Nrdg2 protein has been observed in ALS mice (45), hypomethylation of *NDRG2* in cfDNA of ALS patients is consistent with this finding. Furthermore, hypomethylation of *FITM2* in cfDNA of ALS patients may indicate increased expression of FITM2 protein, which promotes lipid droplet formation, and thus could be contributing to excessive, aberrant accumulation of lipid droplets, as noted upon TDP-43 dysfunction (49) and *FUS* mutation (50). Hence, the ability of cfDNA epigenetic signature analysis to uncover specific genes that correlate with disease progression in this small pilot study is a very encouraging result.

We also sought to compare cfDNA epigenetic changes with neurofilaments, since neurofilaments have been suggested as potential biomarkers in ALS. However, because of the small number of patients for whom neurofilament levels were available, we only observed a significant correlation between pNfH CSF levels and ALS disease progression. For this reason, we tested if cfDNA detection of methylation differences in ALS patients correlated with pNfH CSF levels, and we observed significant correlations between methylation changes in two genes, *RIPK1* and *GLP1*. Interestingly, *RIPK1* has been studied extensively in ALS, with one report documenting its up-regulation in human ALS patients and potential utility as a biomarker (43). Because the RIPK1 protein can induce axonal degeneration by promoting inflammation and necroptosis pathways (42, 59), it was nominated as a therapy target, resulting in the development of an oral RIPK1 inhibitor drug as a potential treatment for sALS (https://ClinicalTrials.gov ID: NCT05237284). We examined *RIPK1* expression in the cortex of post-mortem FTD and FTD-ALS patients, and we independently confirmed significant increases in *RIPK1* RNA levels. As a recent single-cell RNA sequencing study of sALS patients identified increased *RIPK1* as a potential factor in promoting glial cell activation and neuroinflammation (51), we examined the function of microglia subjected to RIPK1 expression increase and documented increased inflammatory cytokine production, as well as decreased phagocytosis capacity, increased migration, and elevated cell death, which together indicate that increased RIPK1 expression predisposes microglia to adopt a reactive, proinflammatory state. Detection of the *RIPK1* gene by unbiased cfDNA epigenetic signature analysis underscores the utility of this modality not only for potential biomarker application in ALS, but also for assisting with prioritization of therapy candidates.

In this pilot study, we isolated cfDNA from the blood of ALS patients, carriers, and non-disease controls, and then utilized targeted EM-seq to determine if epigenetic changes could reliably differentiate ALS patients from non-disease controls. We found that the cfDNA epigenetic signatures detected by targeted EM-seq were surprisingly robust, permitting us to accurately diagnose 70% of ALS patients with a blood test. In light of these compelling results, we extended our analysis to test for correlations between cfDNA epigenetic signature in comparison to rate of disease progression and CSF neurofilaments in ALS patients, and we detected sets of genes whose methylation status significantly correlated with ALS-FRS-R decline and with CSF pNfH levels. Given our limited sample size, future studies are needed to determine if cfDNA epigenetic signature analysis using MESA can yield similar results in larger and more diverse ALS patent cohorts. Replication in a much larger cohort is also necessary to rule out the risk of overfitting due to the use of limited sample sizes. Nonetheless, our findings indicate that cfDNA epigenetic signature analysis deserves further evaluation as a potential biomarker in ALS, especially when one considers that the half-life of circulating DNA is exceedingly short, ranging from several minutes to a few hours (13, 54), underscoring its potential utility for effectively and dynamically monitoring effects of experimental treatments in future human clinical trials.

## Methods

### Sex as a biological variable

This study includes analysis of 60 ALS patients and controls, and the study was not powered to evaluate sex as a biological variable. The sex of the study participants is given in Table 1. For experiments using postmortem human tissue, roughly equal numbers of males and females were analyzed, and sex was not evaluated as a biological variable.

### Study cohort

Patients were enrolled at the Department of Neurology, University of Ulm Germany between 2009 – 2023. Patients with ALS were diagnosed with definite, probable, or possible ALS according to revised El Escorial criteria. C9-carrier and C9-ALS were positively tested for a GGGGCC hexanucleotide expansion in the first intron in the C9orf72 gene. Patients classified as sALS tested negative for the 43 most frequent disease-causing mutations and risk factors (Supplemental Table 3). ALS severity was measured using the ALS Functional Rating Scale-Revised (ALS-FRS-R) at time of blood drawing, which represents a quantitative measurement of physical functioning on a scale from 0 (not functional) to 48 (normal function). ALS disease progression was calculated as loss of ALS-FRS-R points from the date of blood drawing compared to the date of first onset of symptoms. The control cohort consisted of patients with cranial nerve palsy (N. III, N. VI, or N. VII palsy), vestibular neuronitis, or benign paroxysmal positional vertigo (BPPV). All participants were subject to a neurological examination and cranial imaging using MRI. Blood samples were obtained from participants recruited between 2009 – 2023 from the Department of Neurology at the University Hospital Ulm via participation in the MND-NET cohort study.

### Collection and preparation of samples

After written consent, whole blood was collected from patients at the ALS clinic at the University Hospital in Ulm. Instead of plasma, we used blood serum for isolation of cfDNA in this study, because serum had been stored at -80°C at the University Hospital Ulm biobank. We used 1.5 – 2.5 mL blood serum as input material for cell-free DNA isolation using the nRichDx Revolution Max20 cfDNA Isolation Kit (PN 100131) according to the manufacturer’s instructions. To rule out significant genomic DNA contamination, we examined isolated cfDNA on an Agilent Bioanalyzer and no large genomic fragments were observed (Figure 1). cfDNA was then shipped on dry ice to TwistBioscience^®^ for methylome sequencing. The quality and quantity of extracted cfDNA was assessed by TwistBioscience^®^.

### Targeted EM-seq of cfDNA

A total of 5 ng cfDNA along with 0.2 pg of unmethylated Lambda DNA per specimen was used to prepare the barcoded NGS libraries using the NEB Next Enzymatic Methyl-seq Kit (New England Biolabs, USA) according to the manufacturer’s instructions. The captured libraries were then supplemented with a 20% PhiX genomic DNA library to increase base calling and submitted to TWIST for Illumina 150 bp paired-end sequencing with ∼45x coverage. The libraries were then hybridized with the Twist Human Methylome Panel that targets 3.98M CpG sites through 123 Mb of genomic content. The panel detects the most current, annotated, and biologically relevant CpG methylation regions in the genome including promoters, enhancers, CpG islands, shores, and shelves. Unmethylated lambda DNA will be used as a control to monitor the C-to-T conversion efficiency. Targeted EM-seq increases detection sensitivity while decreasing sequencing costs (14). Using this approach, we achieved ∼45x coverage, while traditional whole-genome bisulfite sequencing typically achieves only 30x coverage. After removing low-quality reads and adaptor contamination with TrimGalore (https://github.com/FelixKrueger/TrimGalore), we aligned remaining reads to the hg38 human genome reference using bwa-meth (v0.2.7; https://github.com/brentp/bwa-meth), Samtools (v1.6) (60), and Sambamba (v1.0.0) (61). We further remove duplicated reads with Picard Toolkit (v3.0.0; https://broadinstitute.github.io/picard/).

### Differential methylation analysis

For differentially methylated CpGs (DMCs), after removing CpG sites with > 10% missing values for Control vs. sALS patients/C9-carrier comparison, we calculated p-values for each CpG site Mann-Whitney U-test. For de-novo differentially methylated regions (DMRs) discovery, we applied metilene (v0.2.8) (62), and subsequently recalculated Mann-Whitney U-test p-values by combining methylation levels of CpG sites covered in each DMRs. The p-values were adjusted for multiple testing using the Benjamini–Hochberg procedure (https://github.com/statsmodels/statsmodels).

### Deconvolution analysis and in silico simulation

The tissue-of-origin of cfDNA samples were estimated using the UXM deconvolution algorithm (20). We grouped 39 different cell types in the deconvolution reference into seven broad tissue categories: blood immune cells, connective tissue, epithelium, sex-related tissue, neural tissue, muscle, and other. For brain cell subtype deconvolution, an expanded cell-type reference was generated from DNA methylation atlas (20) and single-cell DNA methylation samples (21), using ‘uxm build’. For in silico simulation of UXM, we included each of 5 neuron samples from the DNA methylation atlas (20), then iteratively reconstructed the cell-type reference excluding the held-out neuron sample, randomly mixed the sample into background leukocyte sequencing reads with wgbstools (20) for 50 times, then applied the same UXM deconvolution pipeline to infer cell-type proportion in each mixture.

### Multimodal epigenetic feature extraction

From cfDNA sequenced with targeted EM-seq and whole-genome bisulfite sequencing data obtained from an independent cohort (40), we extracted four types of epigenetic features: cfDNA CpG island (CGI) region methylation, cfDNA DNase hypersensitive site (DHS) region methylation, nucleosome occupancy, and window protection score (WPS). cfDNA methylation for single CpG sites was calculated with MethylDackel (v0.5.1)[ https://github.com/dpryan79/MethylDackel] on aligned bam files. CGI and DHS region annotations were download for hg38 reference genome from UCSC genome browser, then regional methylation was calculated by averaging CpG site methylation within regions. Nucleosome occupancy was calculated using DNAPOS3 (63) for each of the 1 kb transcription start sites (TSS) and polyadenylation sites (PAS) and averaged with bigWigAverageOverBed (https://github.com/ENCODE-DCC/kentUtils/blob/master/bin/linux.x86_64/bigWigAverageOverBed). As described in (18), average WPS was calculated for each targeted sequencing region.

### Multimodal machine learning model for ALS detection

We applied the multimodal epigenetic sequencing analysis (MESA)(39) to build a multimodal machine learning model for sALS detection. For each of the four modalities (i.e. CGI methylation, DHS methylation, nucleosome occupancy and WPS), MESA selected a subset of informatic features (respectively 3,000, 2,500, 200, and 200) based on the Mann-Whitney U-test, and then further selected the most representative features as markers (respectively the top 100, 100, 20, and 80) using the Boruta algorithm. A base estimator would be trained based on the selected features. In detail, in each iteration of leave-one-out cross-validation (LOOCV), feature selection and classifier training were performed for each modality within the training set, then a meta-classifier was trained and made predictions for a test set based on the probabilities predicted by each of the four single modality base estimators. For validation of an independent cohort, we trained MESA on 40 or our samples (sALS, n=19; control, n=21) and calculated the predictive performance with the trained model on the independent cohort (sALS, n=10; control, n=8).

### Genetic analysis

DNA was extracted from blood leucocytes. Analysis of the C9orf72 repeat length was performed by fragment length analysis and repeat-primed PCR (RP-PCR) (64). Electrophoresis was performed on an ABI PRISM® 3130 Genetic Analyzer (Life Technologies, Foster City, California, USA). The data were analyzed using the Peak Scanner software (Applied Biosystems, Waltham, Massachusetts, USA). Samples with a saw tooth pattern in the RP-PCR were further analyzed using Southern blot (65). Genetic testing was performed at the Institute of Human Genetics of Ulm University.

### Quantitative RT-PCR analysis of post-mortem patient cortex RNA

RNA was extracted from postmortem human frontal cortex tissue using the RNeasy Plus Mini Kit (Qiagen), following the manufacturer’s protocol. A total of 500 ng of high-quality RNA (RNA integrity (RIN) > 8 assessed by the Agilent 2100 Bioanalyzer, Agilent Technologies; and concentration by the NanoDrop spectrophotometer, Thermo Fisher Scientific) were used for reverse transcription using the High-Capacity cDNA Reverse Transcription Kit (Applied Biosystems), according to the manufacturer’s instructions. Quantitative PCR was performed in triplicate reactions using SYBR GreenER qPCR SuperMix (Invitrogen) on a QuantStudio™ 7 Flex Real-Time PCR System (Applied Biosystems). Interassay control samples were included on each plate to correct for potential inter-plate variability. For each well of a 384-well plate, 2 µl of diluted cDNA was combined with 2.5 µL SYBR GreenER SuperMix and 0.5 µL of a primer mixture containing 1 µM forward primer and 1 µM reverse primer at a final concentration of 0.1 µM per primer. The following primers were used: *GAPDH* F-GTTCGACAGTCAGCCGCATC, *GAPDH* R-GGAATTTGCCATGGGTGGA; *RPLP0* F′-TCTACAACCCTGAAGTGCTTGAT; *RPLP0* R-CAATCTGCAGACAGACACTGG; *NDRG2* F-CCCGGACACTGTTGAAGG; *NDRG2* R-GGCTGAAAAGATGTCCAAGG; *RIPK1* F-GTACCTTCAAGCCGGTCAAA; *RIPK1* R-TTACTCTGCAGGCTGGGC.

### Studies of iTF-microglia

iTF-iPSCs were differentiated glia using a previously established protocol from the indicated iPSC cell line (52). RNA was isolated using Zymo Quick-RNA Miniprep Kit (Cat #R1055). RNA was reverse transcribed into cDNA using qScript™ Ultra SuperMix (VWR, Cat #76533-178). IL-6, TNF-α, IL-1β, CCL3, and IL-12A gene expression was analyzed using PowerUp™ SYBR™ Green Master Mix for qPCR (Thermo, Cat #A25777) with the following primer sequences: *GAPDH* F-TCCTGTTCGACAGTCAGCCG, *GAPDH* R-CCCCATGGTGTCTGAGCGAT, *IL-6* F-TTCGGTACATCCTCGACGGC, *IL-6* R-TCACCAGGCAAGTCTCCTCA, *TNF-α* F-TGCACTTTGGAGTGATCGGC, *TNF-α* R-CTCAGCTTGAGGGTTTGCTACA, *IL-1β* F-CGAGGCACAAGGCACAACAG, *IL-1β* R-GGTCCTGGAAGGAGCACTTCA, *CCL3* F-GGCTCTCTGCAACCAGTTCTCT, *CCL3* R-GGCTTCGCTTGGTTAGGAAGATGA, *IL-12A* F-CCAGCACAGTGGAGGCCTGTTTA, *IL-12A* R-GGCCAGGCAACTCCCATTAGTTAT.

#### Phagocytosis

We incubated iTF-microglia with pHrodo™ Red Zymosan BioParticles™ Conjugate for Phagocytosis (Thermo, Cat #P35364) at a 1:250 dilution after reconstitution in 2 mL PBS + DAPI. After 5 hrs, cells were placed into a Varioskan™ LUX Multimode Microplate Reader. Fluorescence readings were obtained for DAPI (405 nm) and phrodo™ Red Zymosan BioParticles (585nm). Dynamic range was set to automatic, measurement time 100 ms, excitating bandwidth 12 nm. % Ratio was determined as %RATIO = ((phrodo fluorescence/DAPI fluorescence) x 100). DAPI recordings were used to account for normalization/variability across wells.

#### Migration

We transferred iTF-microglia onto plates containing 100 uM Adenosine 5’-diphosphate (Cayman, Cat #16778) in the bottom chamber and plain media in the upper chamber. After 4 hrs, cells were fixed with 4% PFA for 15 min at RT followed by staining with DAPI for 10 min. The transwell insert was then placed onto a plain glass microscope slide and imaged at a wavelength of 405 using a Nikon eclipse Confocal Ti2 to obtain the total cell count based on DAPI stain. The transwell insert was then placed back onto the glass microscope slide to obtain an image of the cells on the bottom chamber of the transwell (migrated cells). % migration was calculated as %migration = ((migrated cell count/ total cell count) x 100).

#### Cell death

We fixed iTF-microglia with 4% PFA for 20 min followed by three washes with PBS. Cells were then incubated with DAPI for 10 min followed by three washes with PBS. Images were obtained using translucent and 405 filters on the ECHO Revolution at a magnification of 10x. Total cell count per field of view was determined based on DAPI counts, and cell death was determined based on amoeboid+blebbing morphology. % cell death was calculated as % dead cells = ((dead cell counts/total cell counts) x 100).

### Statistical analysis

As indicated, for pairwise comparisons, statistical significance was determined by two-tailed Student’s t-test. For multiple comparisons, statistical significance was determined by one-way analysis of variance (ANOVA) with post hoc Tukey test. We employed the non-parametric Kruskal-Wallis test for comparison of the large post-mortem patient data set. The significance level (α) was set at 0.05 for all experiments. Data were compiled and analyzed using Microsoft Excel or Prism 10.0.0 (GraphPad).

### Study Approval

The study was approved by the Ethics Committee at the University of Ulm (application number 19/12; Ulm, Germany). All study participants provided written informed consent.

### Data Availability

Values for all data points in all six main figures and three supplementary figures are reported in the accompanying Supporting Data Values file. Any additional information on underlying data will be made available by the corresponding authors upon request,

## Acknowledgments

The authors wish to thank A. Cheung and C. Morrison for their technical advice and guidance, A. Beer, S. Hübsch, and D. Schattauer for their assistance with ascertainment of samples, and K. Günther for exceptional data management and organizational support.

## Data Availability

All processed data used to generate the results are available at Zenodo (accession #14625318). Raw sequencing reads are available on the European Genome-phenome Archive (EGA) through accession number EGAD50000001808.

## Author contributions

S.M., C.C., W.L. and A.R.L.S. provided the conceptual framework for the study. S.M., C.C., W.R., C.L.B., L.M.T., W.L. and A.R.L.S. designed the experiments. S.M., C.C., M.M.G.G., F.J.A., Z.W., D.S., A.C.W., and J.D. performed the experiments. S.M., C.C., M.M.G.G., F.J.A., Z.W., C.L.B., L.M.T., A.C.W., D.W.D., L.P., M.P., W.L. and A.R.L.S. analyzed the data. S.M., C.C., W.L. and A.R.L.S. wrote the manuscript.

