## Supplementary Tables & Figures for "Multimodal analysis of cell-free DNA identifies epigenetic biomarkers for amyotrophic lateral sclerosis diagnosis and progression"

**Supplemental Table 1. Tissue deconvolution from blood-derived cfDNA**

| Participant | Neural (%) | Muscle (%) | Blood-immune (%) | Connective (%) | Epithelium (%) | Sex-related (%) | Other (%) |
|---|---|---|---|---|---|---|---|
| Control_1 | 3.17 | 4.76 | 39.37 | 5.30 | 16.42 | 8.90 | 22.07 |
| Control_2 | 4.45 | 6.93 | 40.67 | 4.19 | 15.63 | 8.54 | 19.59 |
| Control_3 | 5.26 | 6.89 | 36.00 | 5.77 | 16.69 | 8.58 | 20.82 |
| Control_4 | 3.51 | 5.98 | 46.28 | 6.32 | 9.45 | 7.98 | 20.48 |
| Control_5 | 3.35 | 4.11 | 50.24 | 4.26 | 16.59 | 8.66 | 12.80 |
| Control_6 | 4.78 | 3.96 | 40.74 | 3.51 | 15.30 | 11.44 | 20.26 |
| Control_7 | 3.40 | 3.88 | 46.55 | 7.31 | 13.89 | 7.49 | 17.47 |
| Control_8 | 2.68 | 5.07 | 41.05 | 5.84 | 16.77 | 8.56 | 20.04 |
| Control_9 | 3.13 | 4.43 | 44.46 | 7.46 | 12.68 | 7.52 | 20.33 |
| Control_10 | 2.71 | 5.02 | 45.53 | 6.96 | 14.38 | 9.25 | 16.15 |
| Control_11 | 4.21 | 5.57 | 37.41 | 6.52 | 19.44 | 7.14 | 19.70 |
| Control_12 | 7.06 | 4.20 | 49.17 | 6.42 | 10.33 | 7.23 | 15.60 |
| Control_13 | 3.96 | 2.82 | 46.65 | 6.75 | 13.52 | 9.56 | 16.74 |
| Control_14 | 3.29 | 3.60 | 44.72 | 8.06 | 14.61 | 8.42 | 17.31 |
| Control_15 | 2.24 | 4.65 | 39.92 | 4.38 | 16.24 | 9.54 | 23.03 |
| Control_16 | 3.08 | 3.97 | 55.15 | 4.16 | 11.62 | 7.71 | 14.31 |
| Control_17 | 5.23 | 3.78 | 37.08 | 3.74 | 23.23 | 7.64 | 19.29 |
| Control_18 | 4.42 | 6.55 | 38.97 | 3.59 | 13.60 | 9.23 | 23.64 |
| Control_19 | 6.24 | 6.51 | 46.61 | 5.73 | 11.65 | 8.45 | 14.80 |
| Control_20 | 5.00 | 4.34 | 55.71 | 3.25 | 9.43 | 5.76 | 16.51 |
| Control_51 | 5.41 | 5.15 | 36.31 | 6.98 | 13.11 | 8.94 | 24.10 |
| sALS_21 | 3.61 | 5.09 | 39.73 | 6.30 | 16.27 | 9.85 | 19.15 |
| sALS_22 | 5.14 | 3.42 | 35.70 | 9.12 | 16.86 | 10.50 | 19.27 |
| sALS_23 | 4.92 | 4.19 | 40.89 | 4.91 | 15.88 | 8.89 | 20.32 |
| sALS_24 | 4.82 | 4.16 | 39.86 | 8.08 | 13.57 | 9.31 | 20.20 |
| sALS_25 | 4.73 | 5.08 | 39.37 | 4.83 | 15.30 | 10.29 | 20.40 |
| sALS_26 | 3.98 | 1.91 | 71.11 | 3.02 | 4.54 | 6.78 | 8.67 |
| sALS_27 | 4.96 | 4.87 | 35.82 | 9.31 | 10.97 | 7.56 | 26.50 |
| sALS_28 | 6.81 | 4.16 | 40.76 | 6.10 | 15.59 | 9.16 | 17.41 |
| sALS_29 | 4.15 | 4.14 | 40.23 | 5.29 | 14.54 | 11.91 | 19.74 |
| sALS_30 | 5.76 | 6.12 | 38.03 | 4.64 | 13.85 | 10.38 | 21.22 |
| sALS_31 | 3.06 | 2.69 | 49.39 | 9.19 | 13.02 | 6.59 | 16.06 |
| sALS_33 | 3.65 | 3.15 | 51.20 | 8.81 | 9.66 | 7.30 | 16.24 |
| sALS_34 | 5.40 | 4.30 | 38.85 | 7.30 | 13.74 | 7.10 | 23.31 |
| sALS_35 | 1.32 | 5.21 | 48.16 | 6.02 | 9.37 | 10.13 | 19.79 |
| sALS_36 | 1.38 | 5.14 | 42.83 | 6.69 | 15.66 | 9.47 | 18.84 |
| sALS_37 | 4.71 | 3.88 | 51.33 | 5.30 | 11.10 | 9.67 | 14.01 |
| sALS_38 | 3.88 | 4.26 | 41.59 | 6.17 | 16.06 | 10.35 | 17.69 |
| sALS_39 | 3.76 | 4.35 | 40.54 | 6.38 | 15.10 | 9.51 | 20.36 |
| sALS_40 | 2.29 | 4.18 | 42.77 | 8.35 | 14.74 | 7.91 | 19.77 |
| C9-Carrier_41 | 5.74 | 4.29 | 36.69 | 8.15 | 16.03 | 10.09 | 19.01 |

| | | | | | | | |
|---|---|---|---|---|---|---|---|
| C9-Carrier_42 | 9.65 | 4.93 | 41.96 | 5.27 | 12.38 | 8.61 | 17.21 |
| C9-Carrier_43 | 1.35 | 4.57 | 32.31 | 5.60 | 20.94 | 9.81 | 25.43 |
| C9-Carrier_44 | 5.35 | 4.57 | 34.12 | 7.72 | 13.45 | 12.90 | 21.89 |
| C9-Carrier_45 | 3.68 | 5.71 | 44.16 | 4.86 | 11.30 | 10.78 | 19.51 |
| C9-Carrier_46 | 4.61 | 5.79 | 36.81 | 6.06 | 12.97 | 10.43 | 23.33 |
| C9-Carrier_47 | 5.71 | 3.81 | 42.94 | 4.21 | 15.61 | 8.83 | 18.89 |
| C9-Carrier_48 | 5.21 | 4.85 | 37.99 | 5.50 | 15.07 | 11.20 | 20.18 |
| C9-Carrier_49 | 0.90 | 4.65 | 43.37 | 6.44 | 12.93 | 11.18 | 20.53 |
| C9-Carrier_50 | 2.03 | 5.10 | 47.23 | 5.14 | 15.32 | 7.35 | 17.83 |
| C9-ALS_66 | 3.83 | 4.94 | 29.38 | 5.08 | 14.96 | 8.92 | 32.90 |
| C9-ALS_67 | 3.62 | 5.23 | 49.52 | 5.71 | 11.56 | 7.55 | 16.81 |
| C9-ALS_68 | 4.57 | 4.91 | 40.48 | 5.49 | 12.49 | 10.34 | 21.73 |
| C9-ALS_69 | 6.31 | 5.77 | 35.84 | 6.00 | 16.22 | 9.75 | 20.12 |
| C9-ALS_70 | 4.26 | 6.31 | 37.79 | 5.69 | 17.04 | 9.81 | 19.10 |
| C9-ALS_71 | 5.57 | 3.93 | 45.56 | 6.32 | 14.03 | 10.26 | 14.32 |
| C9-ALS_72 | 5.23 | 6.13 | 42.23 | 6.50 | 12.79 | 8.89 | 18.23 |
| C9-ALS_73 | 4.51 | 5.49 | 43.90 | 6.91 | 13.18 | 9.46 | 16.54 |
| C9-ALS_74 | 5.14 | 5.24 | 37.86 | 5.43 | 17.79 | 9.69 | 18.86 |
| C9-ALS_75 | 4.29 | 6.09 | 35.20 | 6.32 | 15.59 | 8.05 | 24.45 |

cfDNA = cell-free DNA

**Supplemental Table 2. Inter-individual biomarker variability of the top 100 differentially methylated CpG sites**

| Top N | All | Control | sALS | All | Control | C9-ALS | All | Control | C9-Carrier |
|---|---|---|---|---|---|---|---|---|---|
| 50 | 0.630 | 0.619 | 0.621 | 0.250 | 0.117 | 0.160 | 0.592 | 0.488 | 0.750 |
| 100 | 0.591 | 0.572 | 0.588 | 0.242 | 0.127 | 0.184 | 0.559 | 0.437 | 0.771 |
| 200 | 0.552 | 0.530 | 0.555 | 0.273 | 0.177 | 0.231 | 0.531 | 0.439 | 0.656 |
| 500 | 0.482 | 0.468 | 0.468 | 0.307 | 0.237 | 0.279 | 0.493 | 0.421 | 0.579 |
| 1000 | 0.457 | 0.449 | 0.433 | 0.331 | 0.271 | 0.319 | 0.469 | 0.405 | 0.538 |
| 2000 | 0.410 | 0.405 | 0.383 | 0.347 | 0.298 | 0.337 | 0.431 | 0.367 | 0.502 |

sALS = sporadic ALS patients, C9-ALS = patients with a repeat expansion in the C9orf72 gene, C9-Carrier = asymptomatic carrier of a repeat expansion in the C9orf72 gene

**Supplemental Table 3. Tested ALS mutations and risk factors**

| |
|---|
| ALS2, ANG, ARHGEF28, ATXN2, BSCL2, C9orf72, CCNF, CHCHD10, CHMP2B, DCTN1, ERBB4, FIG4, FUS, GBE1, GLE1, GRN, HNRNPA1, HNRNPA2B1, HSPB1, HSPB8, MAPT, MATR3, MME, NEFH, NEK1, OPTN, PFN1, PRPH, SETX, SIGMAR1, SOD1, SPG11, SPG20, SQSTM1, TAF15, TARDBP, TBK1, TUBA4A, UBQLN2, VAPB, VCP, VEGFA, VPS54 |

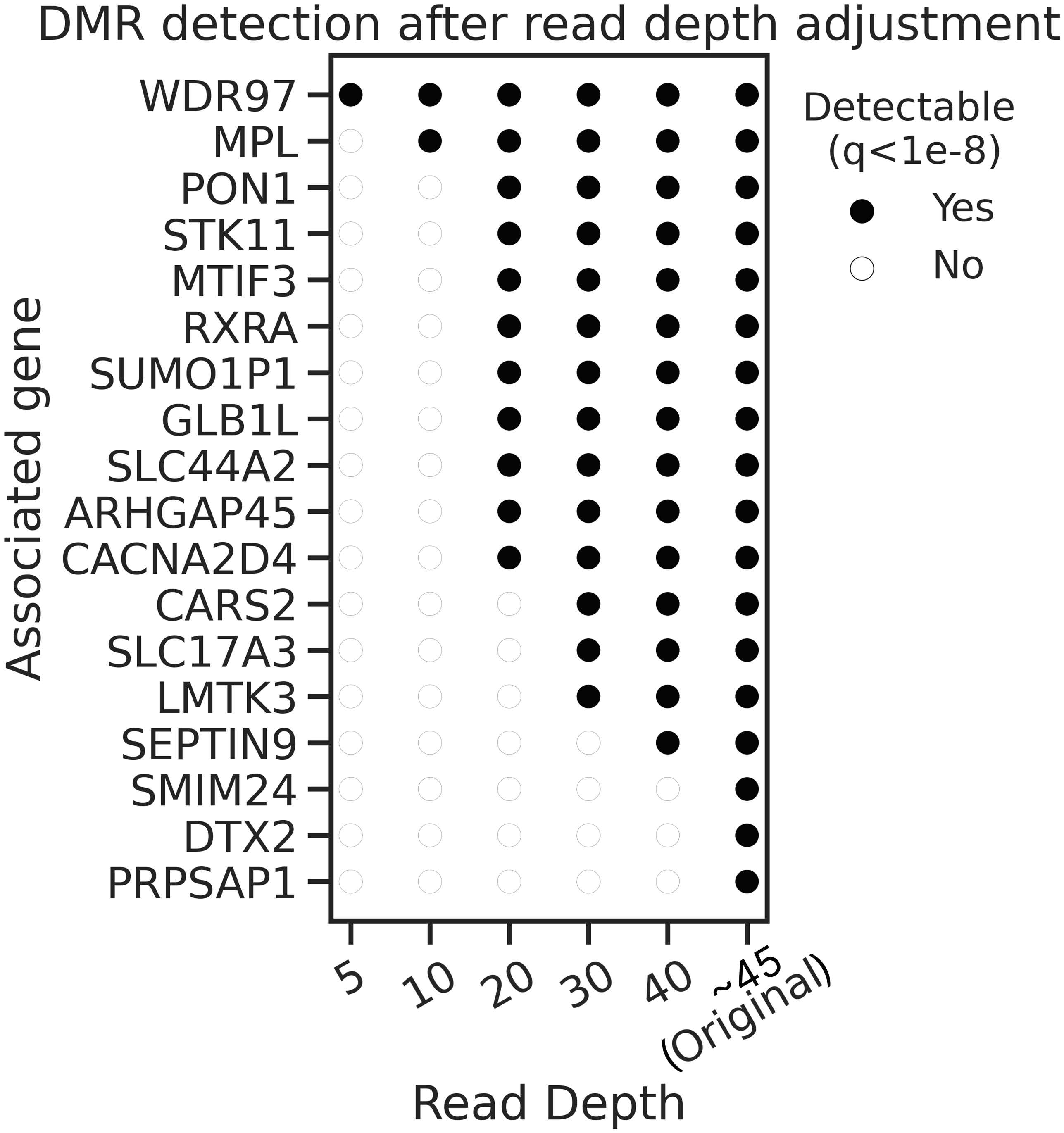


**Figure S1. Extent of read depth required for successful detection of differentially methylated regions in epigenetic sequencing analysis of cell-free DNA**

Detection of differentially methylated regions (DMRs) at q < 1e-8 across varying read depths (X-axis) for different specific genomic regions (Y-axis). The Y-axis represents DMR-associated genes identified from our original analysis.

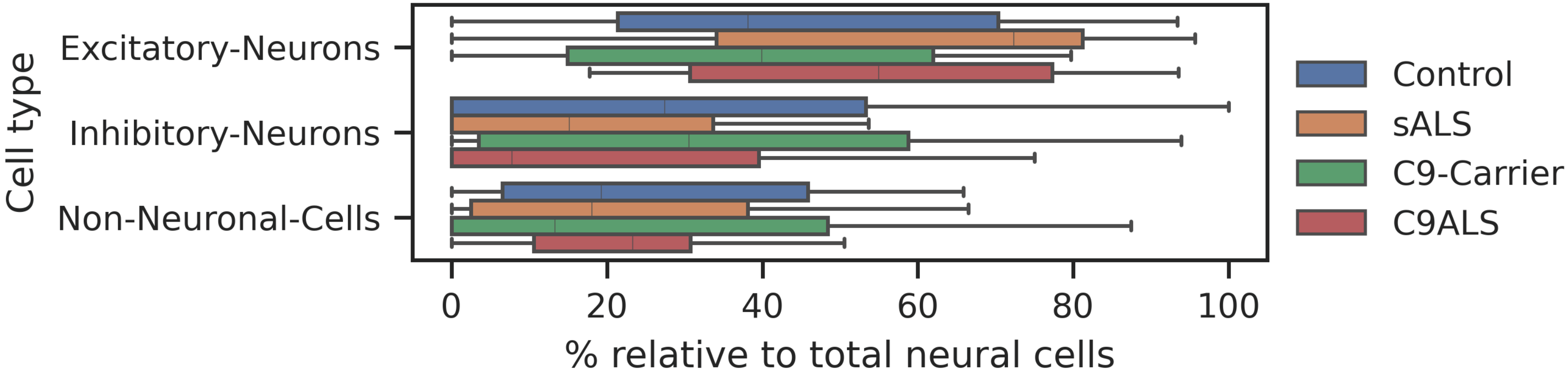


**Figure S2. Determination of specific cell type contribution to the neural component of cell-free DNA**

UXM deconvolution analysis using an extended reference atlas of 188 brain cell subtypes. Boxplots show that three brain cell types emerge as the major contributors to the neural cell-free DNA signal, with the relative contribution to total neural cells of each major contributing cell type indicated.

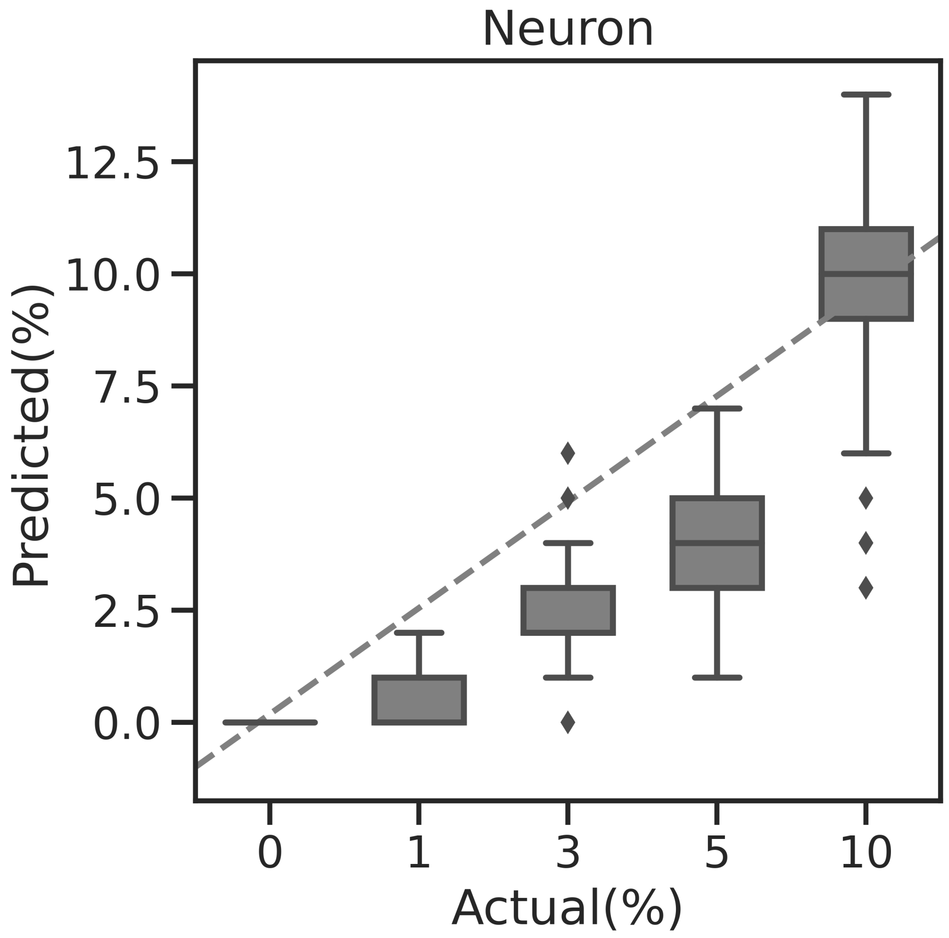


**Figure S3. Validation of UXM deconvolution algorithm for determination of tissue type representation for cell-free DNA sample analysis**

UXM deconvolution analysis of 50 mixtures of five different neuron samples, where the relative concentration of neuron input ranged from 0 to 10%. Here we see the UXM deconvolution algorithm result shown as ‘Predicted %’ on the Y axis plotted against the ‘Actual %’ concentration of neuron input to the various mixtures. The UXM algorithm was applied to all cell-free DNA samples used in this study to determine their tissue-type composition, which is shown in Figure 2C.
